# Circular mRNA encoded PROTAC (RiboPROTAC) as a new platform for the degradation of intracellular therapeutic targets

**DOI:** 10.1101/2022.04.22.489232

**Authors:** Jiali Yang, Jiaojiao Sun, Jiafeng Zhu, Yaran Du, Yiling Tan, Lixiang Wei, Yang Zhao, Qiangbo Hou, Yan Zhang, Zhenhua Sun, Chijian Zuo

## Abstract

Although proteolysis targeting chimera (PROTAC) technology that hijacking the ubiquitin-proteasome system (UPS) to artificially degrade protein is an emerging promising technology for many undruggable targets, it still faces several challenges, such as the difficulty to select high specificity molecule to protein of interest (POI), and only few of the E3-ligase are suitable for PROTAC mediated protein degradation. Protein-based PROTAC, termed BioPROTAC, owns the advantage of specificity but lacks membrane permeability. Here, we develop a new targeted protein degrading platform, which we termed RiboPROTAC, by encoding BioPROTAC protein degraders with circular mRNA (cmRNA) and delivering intracellularly. In this way, intracellular protein GFP and nuclear protein PCNA were successfully degraded. For PCNA targeting, not only the level of target protein is strongly decreased, but also the cell function related is strongly interfered, such as cell proliferation, apoptosis, as well as the cell cycle. What is more, for the first time, we observed that degrading PCNA with RiboPROTAC causes immunogenic cell death (ICD), although not strong, but was verified in vivo. Importantly, our in vivo results demonstrated that cmRNA encoded PCNA degrader exhibited significant tumor suppression effect, thus we provide the first proof of concept for the application of RiboPROTAC as potential mRNA therapeutic. We consider that RiboPROTAC is a new and superior PROTAC technology for targeting the undruggable targets.

## Introduction

Since its first proof of concept reported in 2001, PROTAC technology has greatly developed in scientific research and drug development (1). The fundamental components of PROTAC contain ligand that binds POI, flexible linker and ligand that recruits E3 ligase. The target protein is eliminated though UPS via catalytic-like process. Compared to RNA interference (RNAi) which targets mRNA or clustered regularly interspaced short palindromic repeats (CRISPR) which targets genomic DNA, PROTAC induces rapid and strong protein degradation no matter the half-life of target protein (2). Due to its event-driven but not occupancy-driven mechanism (3), a lot of undruggable targets become druggable, such as transcription factors (4). There are several advantages of canonical PROTAC, such as its catalytic mechanism of action, small-molecule nature, temporary control, and portability (5). However, there are invisible disadvantages as well, such as long discovery phase, off-target effects, hook effect and requirements for ligandable molecules (6). To overcome the notable shortage of low target binding specificity that mediated by small molecule ligand, BioPROTAC was developed (7). Compared to small molecules, the warheads of BioPROTAC are based on protein-binding peptide, DARPins or nanobody thus preserving higher specificity, lower off-target effect and shorter discovery phase. What’s more, BioPROTAC contains E3 ligase itself, thereof it doesn’t rely on the natural expression of endogenous E3 ligase and thus would not break the balance between the E3 ligase and its intrinsic target proteins (7). However, the application of protein-based BioPROTAC is limited by its difficulty of intracellular delivery. So far, its application is mainly focusing on biological discoveries since its delivery relies on protein electroporation, genetic engineering or cell penetrating peptides, which is not always applicable. mRNA technology is revolutionizing drug development nowadays, especially in the development of vaccines and protein replacement therapeutics (8). We proposed that BioPROTAC proteins could be encoded and delivered intracellularly by mRNA whose intracellular delivery has been extensively studied and applied in clinical market successfully. In our previous studies, we applied our circular mRNA, termed cmRNA, to express various kinds of proteins with high and durable protein expression, and successfully used in vivo as anti-tumor therapeutic medicine in preclinical mouse models (9). Here, we demonstrated that cmRNA-encoded BioPROTAC, which we termed RiboPROTAC, is a new platform for intracellular target protein degradation, and for the first time, we successfully applied it in vivo as potential anti-tumor therapeutic by degrading the tumor target PCNA.

## Results

### Validation of RiboPROTAC for degrading H2B-GFP

As illustrated in the scheme in Fig. 1A, the RiboPROTAC cmRNA encodes proteolysis targeting chimera containing a binding domain, a flexible linker and an E3 ligase. Once delivered into cells, RiboPROTAC is translated and followed by target protein ubiquitination and degradation through proteasome pathway. As a proof of concept, green fluorescence protein (GFP) was chosen as the model protein of POI since its convenience for monitoring. On account of the lack of ubiquitination sites in original GFP protein, histone 2B tagged GFP (H2B-GFP) was used instead as the protein degradation target, and the corresponding BioPROTAC Darpin-GFP-SPOP167–374 (DASP) was used according to the report from Lim, et. al (7). Firstly, we transfected HEK293T with different quantities of cmRNA encoding H2B-GFP, and 6 h later these cells were transfected with different quantities of cmRNA encoding GFP degrader DASP. As shown by fluorescence imaging results (Fig.1B), the H2B-GFP was efficiently degraded by GFP degrader. The higher concentration of GFP-degrading mRNA was transfected, the more clearance of signal was observed, with the highest clearance rate as 75% (Fig.1B). To prove that cmRNA mediated GFP degradation is through proteasome pathway, different concentrations of proteasome inhibitor MG132 was added into cells that transfected with cmRNA encoding GFP and/or GFP degrader. As shown by Fig.1C, the GFP clearance rate significantly decreased with increasing MG132 concentration. The GFP clearance was almost abrogated when MG132 concentration reaches 20 μM, demonstrating that proteasome plays a key role in RiboPROTAC mediated GFP degradation.

**Figure.1.**
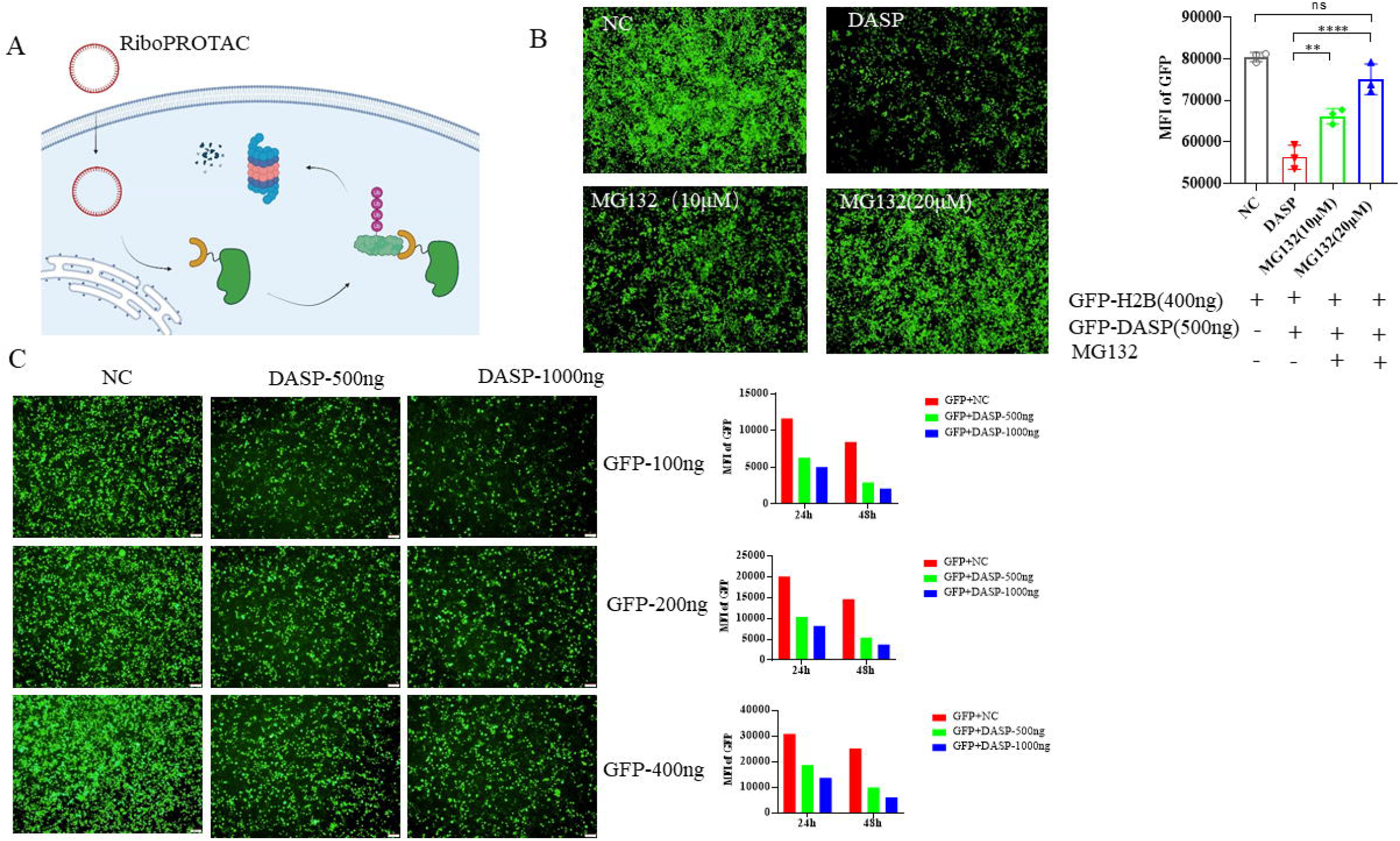
In vitro validation of RiboPROTAC by targeting GFP. (A) Scheme of RiboPROTAC; (B) cmRNA encoded H2B-GFP was degraded by GFP RiboPROTAC, and the degradation was reversed by 10 μM and 20 μM MG132; (C) 100, 200 and 400 ng of cmRNA encoding H2B-GFP was transfected to 293 T, and H2B-GFP was degraded by GFP RiboPROTAC in a dose-dependent manner.

### PCNA degradation mediated by RiboPROTAC

To extend the application of RiboPROTAC platform, we evaluated its capacity of degrading endogenous nuclear protein, PCNA. cmRNA encoding BioPROTAC that targets PCNA was generated by in vitro transcription (IVT), circularization and purification. The PCNA BioPROTAC degrader, named PCNA-SP, which consists of a PCNA binding peptide, a flexible linker and the E3 ligase SPOP167–374, is from the report of Lim et.al (7). Owning to the high homology of human and mouse PCNA sequences, and the peptide-binding site is highly conserved, we evaluated the PCNA degradation effect of RiboPROTAC both in human and mouse cells. As a negative control, the loss of function mutated counterpart of PCNA degrader, named PCNA-SP-Mu, contains a loss of function PCNA-binding peptide. Western blot results showed that the PCNA clearance was as high as 90%, as shown in Fig.2A. Moreover, we found that the cell function was strongly influenced by the elimination of PCNA protein. Firstly, the cell proliferation of cancer cell line A549 and B16F10 were both extremely inhibited upon PCNA protein degradation, as indicated in Fig.2B. Moreover, the AnnexinV/PI assay was performed 24 h post transfection, and the early apoptosis rate of PCNA degraded cells was one-fold higher in A549 and two folds higher in B16F10 cells than control group, as featured by AnnexinV+/7AAD-staining (Fig.2C). The cell cycle analysis revealed that cells were arrested in G1/G0 phase, as illustrated by Fig.2D. In a word, PCNA degradation by RiboPROTAC arrested cells in G1/G0 phase, inhibited their proliferation and promoted cell death.

**Figure.2.**
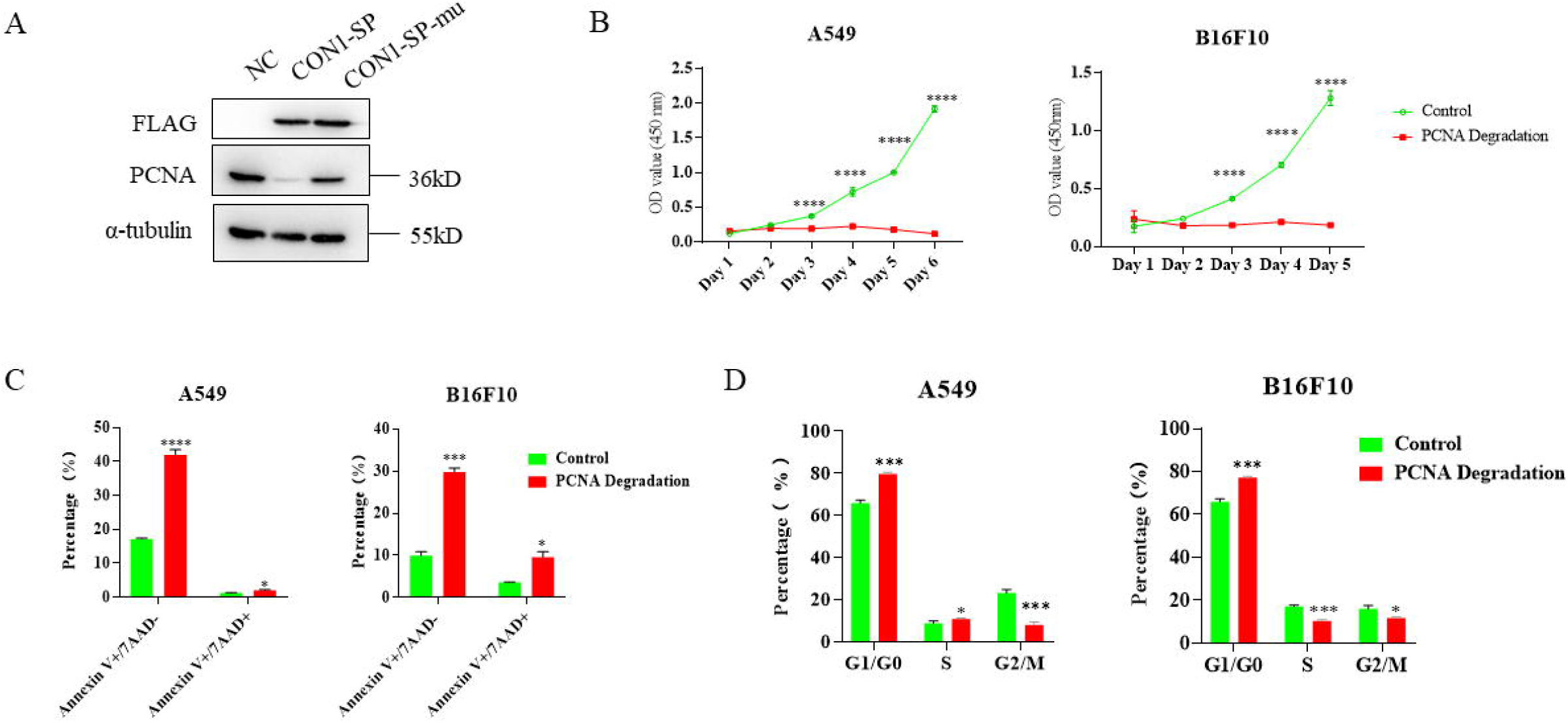
In vitro validation of PCNA protein degradation by PCNA RiboPROTAC, and the resultant cell regulations. (A) endogenous PCNA protein was degraded by RiboPROTAC that encodes FLAG-tagged PCNA degrader CON1-SP, however, was not affected by the loss of mutation counterpart CON1-SP-Mu; (B) cell proliferation of A549 and B16F10 cell line were significantly inhibited by PCNA degradation; (C) Annexin V/7AAD assay results indicated PCNA RiboPROTAC induced notable cell apoptosis of A549 and B16F10; (D) PCNA RiboPROTAC induced cell cycle arrested in G1/G0 phase.

### Degrade PCNA induces slight ICD signature

ICD is a critical event in tumor immunotherapy by triggering the activation of anti-tumor immunity, and facilitates the tumor suppression effect (10). Cell death can induce immune tolerance or immunogenicity, depending on whether a release of damage-associated molecular patterns (DAMPs) occurs during cell death (11). It is still unknown whether PCNA blockade results in ICD. Since we observed strong apoptosis effect in PCNA degraded cell, we tried to investigate if there is ICD involved in PCNA degradation mediated cell deaths. To validate this, we first measured canonical markers from the release of ICD cells, including calreticulin (CRT) and ATP. In our study, a slight but not strong ICD signature was observed. For instance, there was higher secretion of ATP in PCNA-degraded B16 cells while not PCNA-degraded A549 cells, as shown in Fig.3A. CRT exposure on the cell membrane was a lot higher on PCNA-degraded A549 cells than on PCNA-degraded B16 cells, as indicated in Fig.3B. Thus, we performed an in vivo assay to evaluate the immunogenicity of cell death induced by RiboPROTAC targeting PCNA. Equal amount of either non-transfected B16 cells, control mRNA-transfected B16 cells or PCNA-degrader mRNA transfected B16 cells were i.t. (intratumoral injection) injected into B16 tumors and the tumor volume was monitored every three days, as illustrated by the scheme in Fig.3C. There was no significance of tumor growth rate among groups that i.t. injected with Ringers’ solution, non-transfected B16 cells and control mRNA-transfected B16 cells. However, there was significant tumor growth inhibition of group that i.t. injected with B16 cells that transfected with PCNA-degrader mRNA (Fig.3D), indicating that the cell death caused by RiboPROTAC evokes part of the host immune system and was immunogenic.

**Figure.3.**
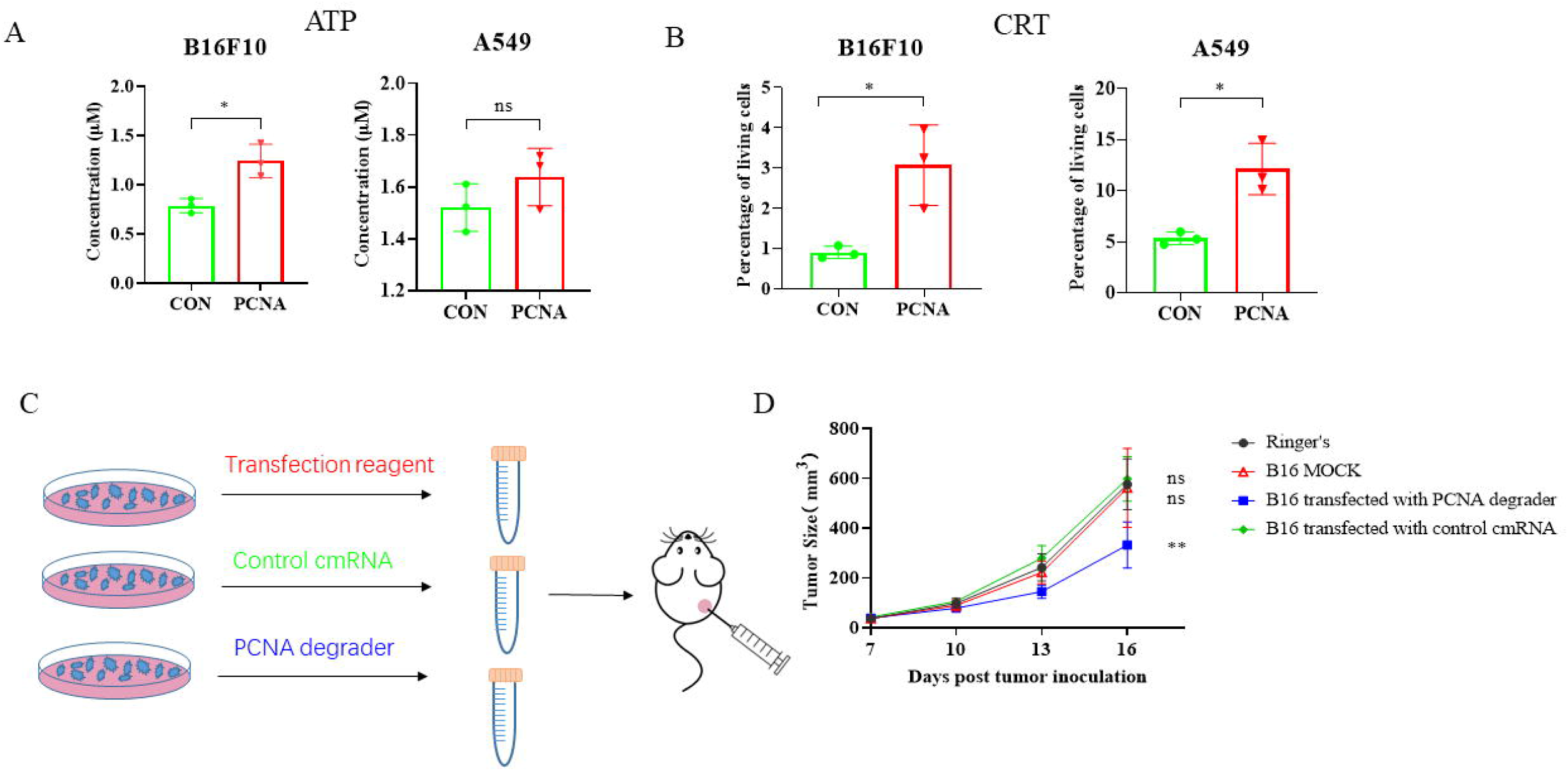
PCNA RiboPROTAC induced slight ICD in B16F10 and A549 cancer cell lines. (A) PCNA RiboPROTAC resulted in significant promotion of ATP release in B16F10, but not in A549; PCNA RiboPROTAC induced elevated CRT level on cell surface of both B16F10 and A549; scheme of the in vivo evaluation of PCNA RiboPROTAC induced ICD; (D) i.t. injection of PCNA RiboPROTAC treated B16F10 cells led to suppression of B16F10 tumor growth in mice.

### Transcriptome profile of PCNA-degradation cells

To date, despite there have been abundant of small molecule PCNA inhibitors developed, none of them are approved for clinical applications. We considered that deeper understanding of the molecular and cellular regulations under PCNA degradation may facilitate the translation of PCNA RiboPROTAC to a clinic therapeutic. To obtain a systematic understanding for the cellular change of mRNA transcription post PCNA degradation, transcriptome profile was generated by transcriptome sequencing and data analysis. As indicated in Fig.4A, there were 758 differential genes with 455 up-regulated and 303 down-regulated (|log2(Foldchange)|>1&Padj<0.05) found in the profile. The heatmap results showed the first 20 down-regulated genes (Fig.4B). And the GO results showed that the differential genes were involved in different, but mostly cell proliferation activities, such as nuclear division, chromosome segregation, mitotic nuclear division, sister chromatid segregation and mitotic sister chromatid segregation (Fig.4C). The KEGG analysis shows that the top 3 pathways that differential genes enriched are cell cycle, DNA replication and p53 signaling pathway (Fig.4D). This gene profile of PCNA degradation is mainly consistent with our previous knowledge of PCNA functions. However, we also found genes involve in signaling pathway that beyond our conventional understanding of PCNA related cell regulations, for example, C-type lectin receptor signaling pathway (shown in Fig.4D), which is important in immunity regulations. The up-regulated genes enriched in C-type lectin receptor signaling include MDM2, TNF, CLEC4E, MAPK11, NFATC4, PLK3, CCL17, CALML5, CCL22 and NLRP3, of which CCL17 and CCL22 may be correlated to the recruitment or activation of immune cells (12-15), and NLRP3 is the gene responsible for the generation of NLRP3 inflammasome, which is important for caspase-1 activation and the secretion of proinflammatory cytokines IL-1β/IL-18 in response to cellular damage (16). Additionally, there is a down-regulation of BIRC5 gene, which is thought to be correlated with tumor microenvironment and immune cells infiltration in a variety of tumors (17).

**Figure.4.**
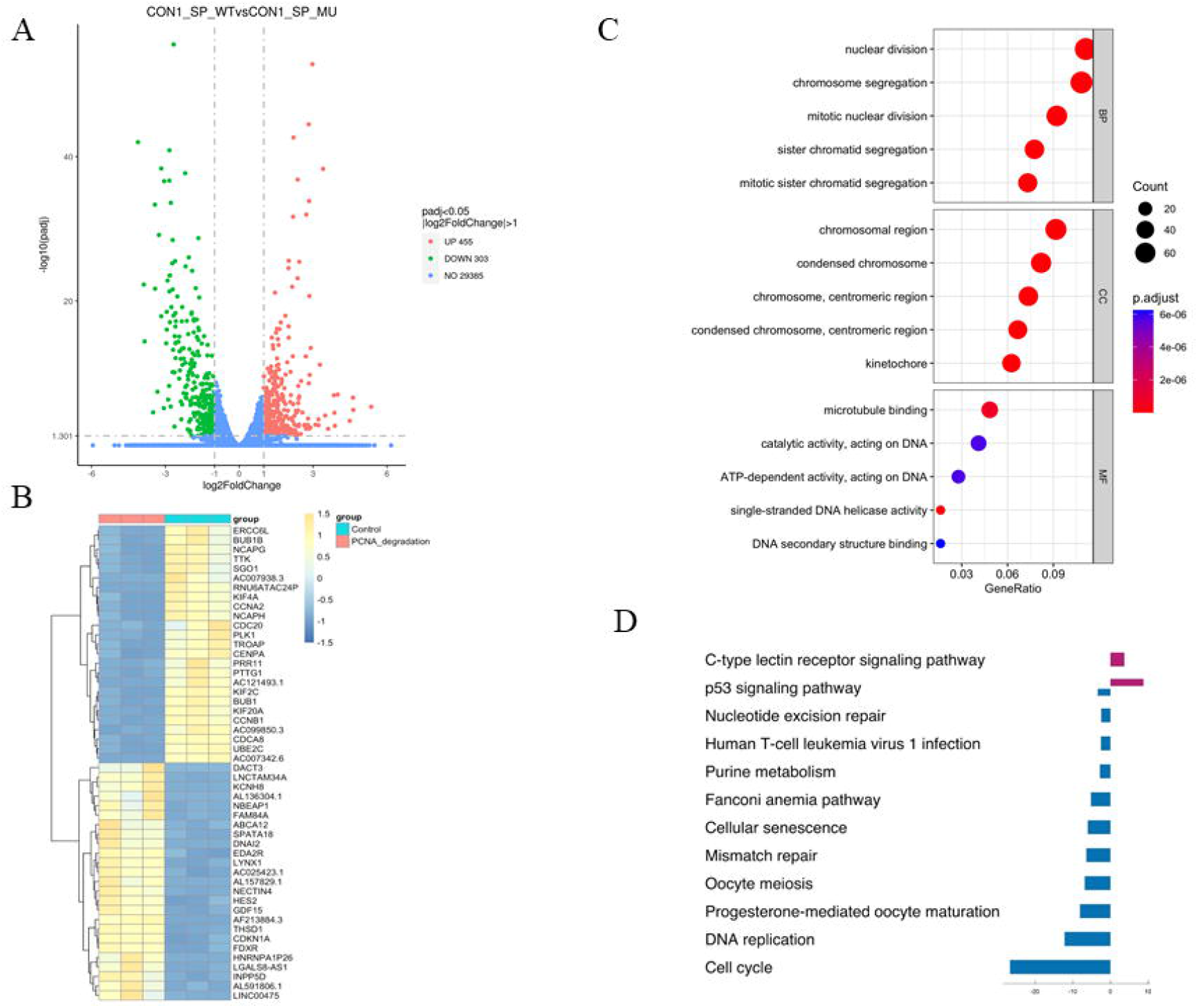
Transcriptome analysis of the B16F10 cells that treated by PCNA RiboPROTAC. (A) 758 differential genes with 455 up-regulated and 303 down-regulated (|log2(Foldchange)|>1&Padj<0.05) were shown; (B) the top 20 up-regulated and down-regulated genes were shown;(C) gene ontology (GO) analysis categorized differential genes to different cell activities; (D) KEGG analysis categorized differential genes to different cell signaling.

### Validation of RiboPROTC application in vivo

The application of protein-based BioPROTAC is limited by its in vivo delivery, which is essential for the development of therapeutics. Since mRNA delivery has been studied extensively, and undergone well-developed (18), we consider that efficient in vivo delivery of cmRNA based RiboPROTAC is highly promising. We have previously shown that direct intratumoral injection of naked cmRNA resulted in durable protein expression in tumor tissues (9). Hence, we used this mouse model to verify RiboPROTAC’s application in vivo. Firstly, we used H2B-GFP as model protein. As illustrated in Fig.5A, when the tumor reaches around 50 mm^3^, we injected 10 μg cmRNA encoding GFP intratumorally and 4 h later, we injected 10 μg cmRNA encoding GFP degrader intratumorally. Tumor were dissected and analyzed by flow cytometry. The percentage of GFP positive cells in control group is around 15%, which indicated that the in vivo intratumoral transfection efficacy of naked cmRNA is about 15% (Fig.5B). Group that injected with cmRNA encoding GFP degrader displays a lower percentage of positive cells, to lower than 5%, and a significant lower level of the median fluorescence intensity (MFI) (Fig.5C). These results prove that, for the first time, RiboPROTAC can be successfully delivered in vivo and applied to degrade target protein.

**Figure.5.**
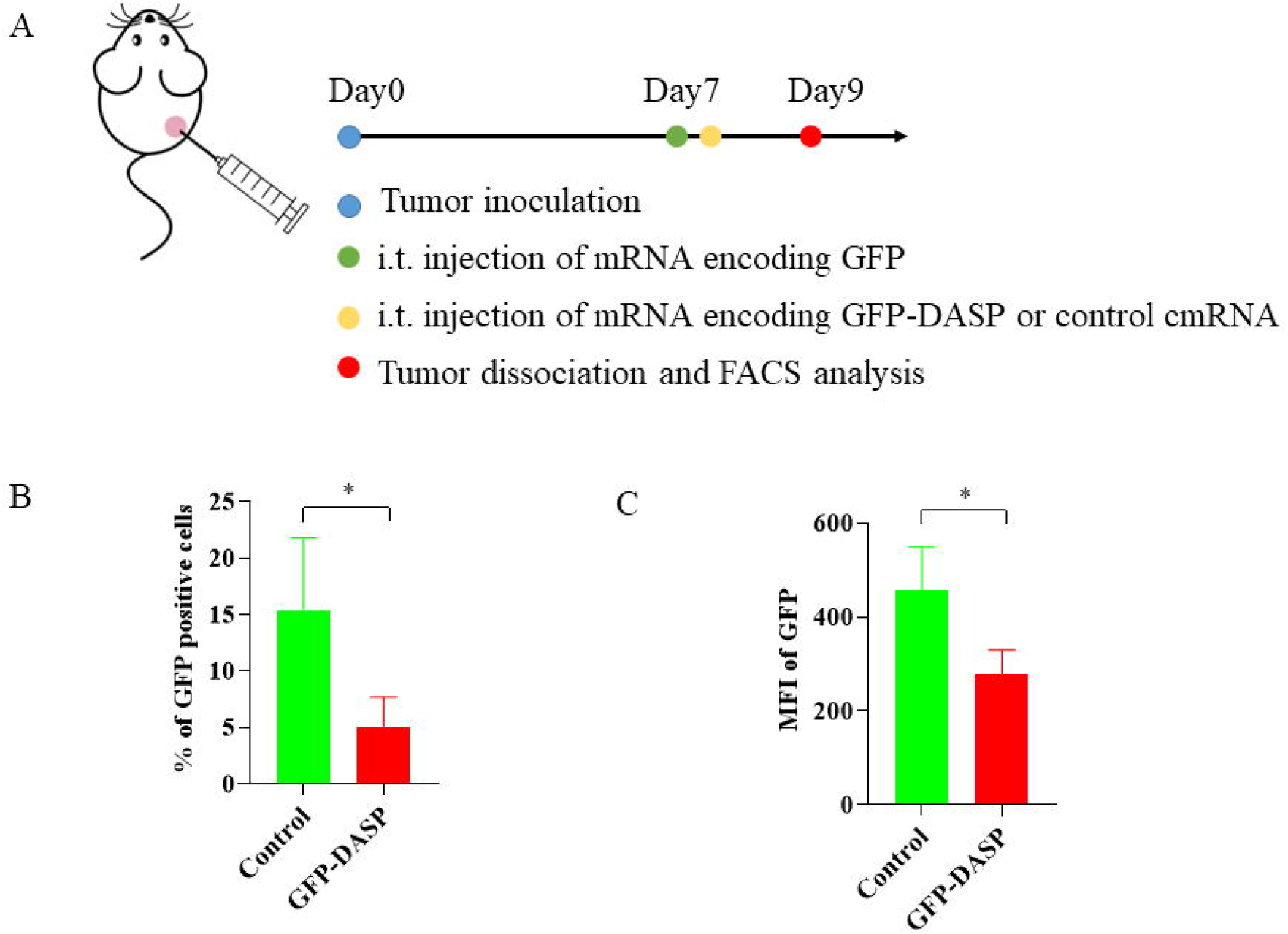
In vivo verification of RiboPROTAC mediated protein degradation using H2B-GFP as model. (A) scheme of i.t. injection of cmRNA encoding H2B-GFP and H2B-GFP degrader, and the expression of GFP in tumor cells was analyzed by FACS; (B) results of FACS analysis revealed reduced percentage of GFP positive cells in H2B-GFP RiboPROTAC group (DASP); (C) results of FACS analysis revealed reduced median fluorescence intensity (MFI) of GFP in H2B-GFP RiboPROTAC group (DASP).

### Antitumor therapeutic effect of RiboPROTAC encoding PCNA degrader

PCNA is overexpressed in many tumor tissues and correlated with the survival of patients, thus is considered as a tumor therapeutic target (19). Encouraged by the in vitro results that PCNA RiboPROTAC induces tumor cell death, as well as the evidence showing that PCNA degradation induces a slight immunogenic cell death signature, moreover, the in vivo delivered cmRNA GFP RiboPROTAC presented GFP protein degradation activity, thus, we prospect that PCNA RiboPROTAC may suppress tumor growth in vivo. To evaluate the antitumor therapeutic effect of PCNA-degrader in vivo, 10 μg of cmRNA encoding PCNA degrader was i.t. injected into mice bearing B16F10 tumors, for negative control, tumors were injected with the buffer Ringer’s solution or PCNA RiboPROTAC loss of function counterpart (PCNA-SP-Mu) (Fig. 6A). First of all, the tumor cell apoptosis was evaluated by TUNEL assay. Fig.6B shows that compared to control group, there was much more tumor cells went apoptosis in PCNA degradation group, however, no significant suppression of tumor growth was observed (data not shown). These results suggest that intratumoral delivered RiboPROTAC induced notable cell death in tumor tissue, despite that the total transfection efficacy of RiboPROTAC naked cmRNA in tumor tissue was rather low, which was revealed by the in vivo GFP degradation assay in Fig.5B. Considering that transfection efficacy by intratumoral injection of naked cmRNA is only around 15%, and PCNA degradation effect is highly correlated with the protein expression of intracellular PCNA degrader, we used transfection reagent in vivojet-PEI® to deliver PCNA degrader in vivo. Empowered by PEI mediated delivery, PCNA RiboPROTAC shows antitumor therapeutic effect in vivo. As illustrated in Fig.6C, tumors that injected intratumorally with PCNA RiboPROTAC that encapsulated with PEI were found to be significantly suppressed, compared to the groups of negative control, single PEI or PCNA mutation RiboPROTAC encapsulated with PEI. These results suggest that cmRNA encoded PCNA degrader is efficient for in vivo tumor therapy in mouse syngeneic tumor model.

**Figure.6.**
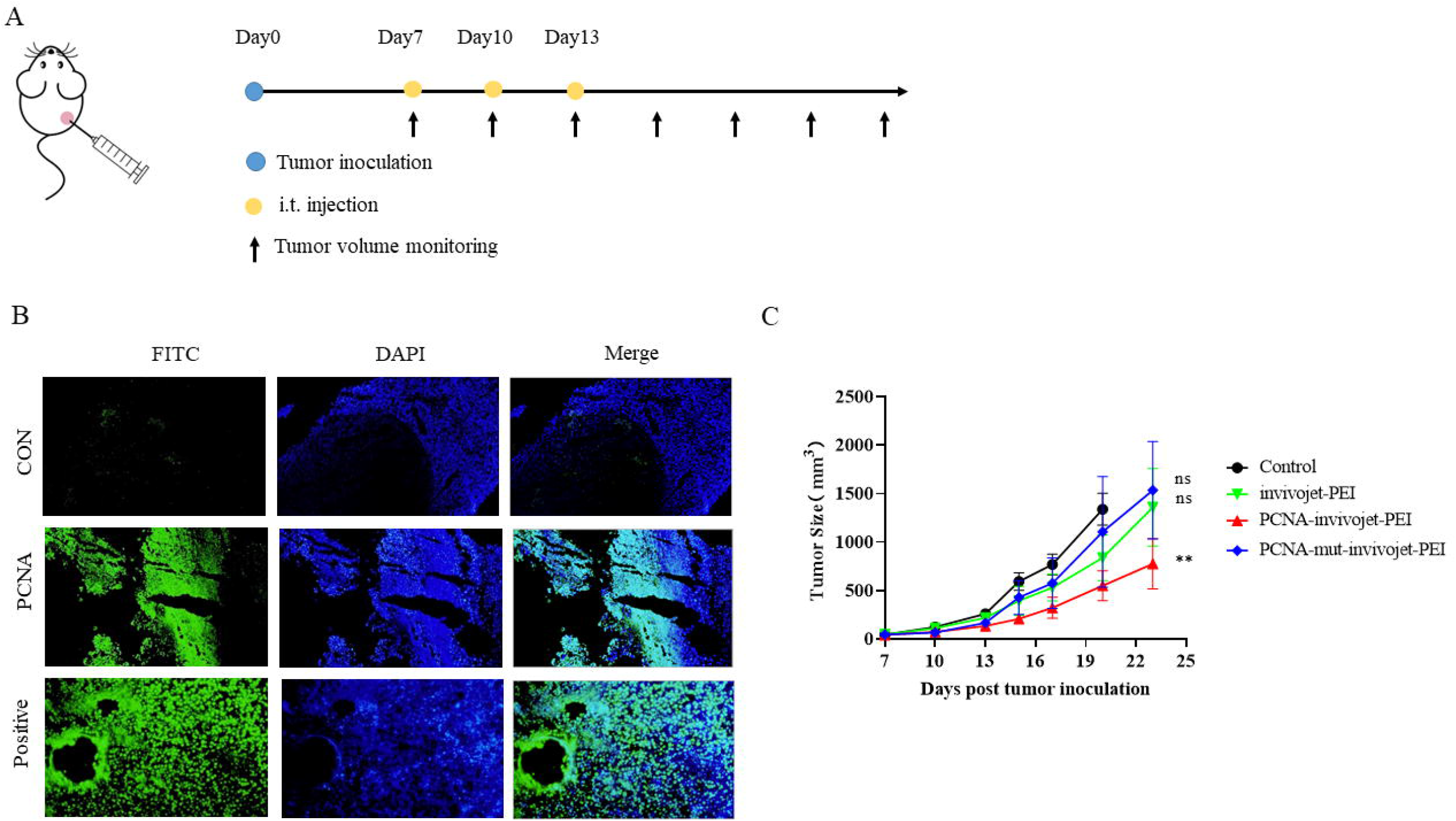
In vivo antitumor effect of PCNA RiboPROTAC. (A) scheme of the i.t. injection of PCNA RiboPROTAC in mice bearing B16F10 tumor; (B) naked cmRNA was i.t injected and TUNEL assay on the section slide of the tumor tissue was performed. Significant cell apoptosis was observed in the group of PCNA RiboPROTAC i.t. B16F10 tumor; (C) mice bearing B16F10 tumor were i.t. injected with Ringers’ solution as control, or singly PEI, or PEI-encapsulated PCNA RiboPROTAC, or PEI-encapsulated loss of function mutation PCNA RiboPROTAC. Only i.t. injection of PEI-encapsulated PCNA RiboPROTAC led to significant suppression of tumor growth.

## Discussion

PROTAC has opened a new area of drug discovery and changed the game of developing small molecule therapeutics through a new mechanism of action. Since its first proof-of-concept, several PROTAC based therapeutic candidates targeting to challenging targets such as androgen receptor, estrogen receptor and Tau protein have shown promising pre-clinical results, and two drug candidates, ARV-110 and ARV-471, have entered Phase II clinical trials (2). Besides the canonical small molecule PROTAC, protein degradation technologies that rely on different mechanisms other than proteolysis were also developed, such as lysosome-targeting chimeras (LYTACs) (20) that dependent on lysosomal protein degradation pathway; autophagy-targeting chimeras (AUTACs) dependent on autophagy pathway (21, 22). There is a dilemma of drug discovery for small molecule-based PROTAC. To increase target range as well as target specificity, it demands large molecular weight and big spatial structure, which in turn hinders the cell permeability and other pharmacokinetic properties. Another potential problem is that limited E3 ligases are available and therapeutic effect is largely dependent on cellular E3 ligase abundance, and mutations of the required E3 ligase may result in drug tolerance. Although sharing the same mechanism of action, BioPROTACs display different pharmaceutical properties and demands different but easier drug design strategies (23). First of all, the warheads of BioPROTAC for POI targeting include peptides derived from natural binding proteins, scFv, nanobody, or DARpins, are considered to possess high specificity for protein targets. In addition, the pre-existence of E3 ligase in BioPROTAC avoids the risk of decreased therapeutic effect by mutation or low abundance of endogenous E3 ligase in cells. And the delivery problem of BioPROTAC could be resolved by gene products such as viruses, DNA or RNA, of which the expression could be fine-tuned by promoter or molecular switches (2).

Our previous results of cmRNA inspired us to further extend its application in target protein degradation through proteolysis. cmRNA which consists of internal ribosome entry site (IRES) element derived from echovirus 29, and improved homology arms and spacers, is a novel circular mRNA platform. In our previous studies, we proved that cmRNA can mediated strong and durable protein expression both in vitro and in vivo. Moreover, our studies demonstrated that cmRNA is sufficient for the expression of various types of proteins (9). Therefore, we prospect that cmRNA is probably an excellent vector to deliver and express BioPROTAC molecules intracellularly. Using H2B-GFP as model protein, we observed as high as 75% of H2B-GFP degradation in cells transfected with RiboPROTAC encoding GFP degrader. Furthermore, the degradation is found largely proteasome pathway dependent, as approved by the results that RiboPROTAC mediated H2B-GFP degradation was abrogated by proteosome inhibitor MG132. Next, to evaluate whether the protein degradation mediated by RiboPROTAC leads to cellular functional change, PCNA was chosen as another target for verification, due to its extremely important role for cell maintenance and proliferation, and its critical role as a tumor therapeutic target. After transfection with PCNA degrader, not only the PCNA protein was found decreased, but also the cell function was strongly interfered, such as a sharp suppression of cell proliferation, high rates of cell apoptosis and significant cell cycle arrest, in cancer cell line B16F10 and A549. These results demonstrated that RiboPROTAC mediated PCNA degradation resulted in potent cytotoxicity for cancer cells in vitro. ICD is considered to exert dual functions in tumor therapy, including a notable reduction of tumor burden by inducing tumor cell death, and a primming of tumor immunity to arouse systemic T cell cytotoxicity for tumor elimination (10). Accumulated pre-clinical and clinical studies have revealed that ICD induction is a crucial and promising strategy in developing anti-tumor therapeutics or clinical therapeutic methods (24, 25). In the current study, for the first time, we observed a slightly immunogenic cell death signature by the detection of in vitro ICD markers. This ICD phenomenon was also verified in vivo by intratumoral injecting B16F10 cell lysates from RiboPROTAC induced cell death. We found that this injection led to a significant suppression of B16F10 tumor growth in mouse syngeneic tumor model, this is probably caused by the injection of released tumor antigens and DAMPs molecules, including CRT and ATP, from RiboPROTAC induced cell lysates, and their synergically induction of tumor immunity. We consider that this is a trustable conclusion, since our in vivo verification method for ICD is concordant to the well-accepted “gold standard” principle for ICD judgement (26). However, it is reasonable to clarify that, ICD induced by PCNA RiboPROTAC may be not sufficient to trigger robust systemic T cell immunity, since the in vitro CRT and ATP release, and the in vivo injection of cell lysate induced tumor suppression are slight.

For a deeper insight into molecular regulations mediated by PCNA degradation, a transcriptome analysis was performed. The results of transcriptome analysis showed that 758 transcripts were changed, with 455 genes up-regulated and 303 genes down-regulated. The KEGG analysis showed that these genes are categorized to pathways involved in cell cycle, DNA replication and p53 signaling, which are in correspondence with the known functions of PCNA. Besides, several genes that associated with tumor immunity are identified, such as CCL17, CCL22 and NLRP3, which are categorized to C-type lectin receptor signaling pathway, and the gene BIRC5 is a regulator for tumor immunity. CCL17 is a chemokine and plays an effective role in chemotaxis of CCR4^+^ T lymphocytes through the CCR4 receptor molecule (27). Previous studies indicated that CCL17 recruits CCR4^+^ lymphocytes to infiltrate tumor and facilitates anti-tumor immune regulations (12). CCL22 is a chemokine that recruits regulatory T cells (Treg) to infiltrate into tumor and contributes to the suppressive immune microenvironment (TME), thus may be a negative regulator for anti-tumor immunity (14, 15). NLRP3 is the key protein for NLRP3 inflammasome (28), which is found to activate the protease caspase-1 to induce gasdermin D-dependent pyroptosis and facilitate the release of IL-1β and IL-18, so as to activate tumor immunity (29). Besides, BIRC5 is found to be correlated with TME regulations and tumor infiltration of immune cells (17). However, the exact role of BIRC5 in tumor immunity remains unclear. Collectively, the changes of gene expression from transcriptome study suggest that CCL17 and NLRP3 may contribute positively to the immunogenicity of PCNA degradation mediated cell death, however, CCL22 may contribute to the suppression of tumor immunogenicity. These complicated gene regulations may explain why PCNA degradation mediated cell death exhibits a slight ICD signature, but not a strong immunity.

Due to the decades of studies of mRNA delivery, there have been significant progressions in this research area recently, and enabled the success of first in human COVID-19 mRNA vaccine in the market (30). Besides, an abundant of mRNA-based vaccines and therapeutics are being evaluated in clinical trials (31), suggesting that the delivery of mRNA in vivo is practicable. Therefore, we considered that, compared to protein-based BioPROTAC, RiboPROTAC owns advantages for in vivo therapeutic applications. Our previous study demonstrated that intratumoral administration of naked cmRNA leads to durable protein expression in syngeneic mouse tumor model (9), in the current study, we evaluated the in vivo delivery and function of RiboPROTAC with the same method. Our results revealed that GFP RiboPROTAC decreased intratumoral GFP protein level, suggesting that intratumoral delivery of naked cmRNA of GFP RiboPROTAC is successful, and targeted GFP degradation occurred efficiently. To the best of our knowledge, this is the first proof of concept experiment to deliver a BioPROTAC molecule in vivo and prove its protein degradation function, in an mRNA way. Using the same method, RiboPROTAC mediated PCNA targeting in vivo was performed. The TUNEL assay results showed an increase of tumor cell apoptosis in PCNA RiboPROTAC group, compared to the untreated one. Importantly, RiboPROTAC power-assisted with delivery reagent invivo-jetPEI showed significant antitumor therapeutic effect in vivo, featuring by a remarkable suppression of B16F10 melanoma tumor growth. No complete response (CR) of the final therapeutic results achieved in this study, this may be attributed to the relative low transfection efficacy of invivo-jetPEI. To the best of our knowledge, this is the first proof of concept experiment to validate that cmRNA encoded RiboPROTACs can be applied in vivo as drug candidates to realize significant therapeutic effects. A variety of carriers for mRNA delivery has been developed and evaluated in clinical studies (31). Among them, lipid nanoparticles (LNPs) and lipoplexes (LPXs) are the two most-developed carriers. LNPs have been applied in the successful development of COVID-19 mRNA vaccines, which are approved by FDA and broadly vaccinated to battle against the SARS-Cov-2 pandemic (30). LNPs are utilized as well for mRNA therapeutics, for instance in the CRISPR-Cas9 mRNA-based gene editing project for the treatment of Transthyretin Amyloidosis, by delivering Cas9 mRNA and gRNA to hepatocytes (32). Thus, we consider that the applications of mRNA delivery by LNP to liver is a promising technology for liver diseases. Recently, LNP was found can be rationally modulated to distribute to specific organs, like liver, lung and spleen, thus RiboPROTAC may be delivered to those organs by LNP (33). The intracellular delivery of RiboPROTAC by LNPs is being investigated in our future studies. LPXs are mainly developed by BioNTech for therapeutic tumor mRNA vaccines and are under evaluation in clinical trials (34). is still to be explored whether LPXs are suitable for intracellular delivery of RiboPROTAC. To sum up, using GFP and PCNA as model targets, we provided a first proof of concept to verified circular mRNA encoded RiboPROTAC as a powerful platform to degrade protein of interest both in vitro and in vivo. We prospect that RiboPROTAC will be an alternative sword to clear the obstacles on the way of drug discovery by targeting numerous undruggable targets, and with efficient mRNA delivery technology, it possesses advantages over the small molecule based PROTAC and recombinant protein based BioPROTAC. The intensive studies and recent progress of mRNA delivery will ensure the in vivo applications and clinical translations of RiboPROTAC in the future.

## Materials and methods

### Gene cloning and vector construction

DNA fragments that containing PIE elements, IRES, coding regions and others were chemically synthesized and cloned into a restriction digestion linearized pUC57 plasmid vector. DNA synthesis and gene cloning were customized ordered from Suzhou Genwitz Co.Ltd. (Suzhou, China).

### cmRNA preparations

cmRNA precursors were synthesized by in vitro transcription from a linearized plasmid DNA template using a PurescribeTM T7 High Yield RNA Synthesis Kit (CureMed, Suzhou, China). After in vitro transcription, reactions were treated with DNase I (CureMed, Suzhou, China) for 15 min. After DNase I treatment, unmodified linear mRNA was column purified using a GeneJET RNA Purification Kit (Thermo Fisher). For cmRNA: RNA was purified, after which GTP was added to a final concentration of 2 mM along with a buffer including magnesium (50 mM Tris-HCl, (pH 8.0), 10 mM MgCl2, 1 mM DTT; Thermo Fisher). RNA was then heated to 55 °C for 15 min, and then column purified. For high-performance liquid chromatography (HPLC), RNA was run through a 30 × 300 mm size exclusion column with particle size of 5 μm and pore size of 1000 □ (Sepax Technologies, Suzhou, China) on an SCG protein purification system (Sepure instruments, Suzhou, China). RNA was run in RNase-free Phosphate buffer (pH:6) at a flow rate of 15 mL/minute. RNA was detected and collected by UV absorbance at 260 nm. Concentrate the purified cmRNA in an ultrafiltration tube, and then replace the phosphate buffer with an RNase-free water.

### Cell lines

Murine melanoma cell line B16F10, human cell line HEK293T and A549 were cultured in DMEM (BI) supplemented with 10% fetal calf serum (Gibco) and penicillin/streptomycin antibiotics (100 U/mL penicillin, 100 ug/mL streptomycin, Gibco). All cell lines were maintained at 37°C, 5% CO2, and 90% relative humidity.

### mRNA in vitro transfection

Cells reaching 60%∼80% confluent were transfected with mRNA using Lipofactamine MessengerMAX (Invitrogen) according to the manufacture’s instruction. Briefly, the mRNA and Lipofactamine Messengermax reagent were diluted with Opti-MEM (Gibco), mixed together and incubated for 5 min at room temperature for complex formation. Then the entire mixture was added to each well and the cells were incubated in the in a 5% CO2 incubator.

### Detections of immunogenic cell death markers

A549 and B16F10 cells were transfected for 24 h and 48h. After incubation, Cells and supernatants were collected separately. The extracellular ATP content was measured with an ENLITEN®ATP Assay system Bioluminescence Detection Kit (Promega) according to the manufacture’s instruction immediately. Cell surface expression of CRT was detected by flow cytometric analysis. The transfected cells were collected with trypsin and cell culture medium, stained using the Anti-hCalreticulin (0.025 μg/100 μl, R&D) and Fluorescein goat anti-mouse IgG (1:100, Invitrogen), analyzed by flow cytometry using AttuneTM NXT.

### Cell-cycle analyses by flow cytometry

A549 and B16F10 cells transfected with CONSP-WT or CONMUSP cmRNA for 24 h were harvested by trypsin, washed with PBS, fixed with 70% ethanol and incubated in -20°C overnight. Then the cells were stained with PI/RNase Staining Buffer (BD Pharmingen) and analyzed by flow cytometry using AttuneTM NXT.

### Cell proliferation assay

We used Cell Counting Kit-8 (Dojindo) to detect the cell proliferation according to the manufacturer’s instructions. After transfected with cmRNA for 24 h, 1000-2000 cells/well were seeded into 96-well plates and cultured for continuous 6 days. 10 μl CCK-8 solution mixed with 90 μl culture medium were added to the cells every 24 h. After 2 h incubation, we used a microplate reader to detect the absorbency at a test wavelength of 450 nm.

### Apoptosis analyses by flow cytometry

A549 and B16F10 cells were transfected with PCNA degradation cmRNA (PCNA-SP) or negative control mRNA (PCNA-SP-Mu) for 24 h. For apoptosis analysis, the transfected cells were collected with trypsin and cell culture medium, stained using the APC Annexin V Apoptosis Detection Kit with 7-AAD (BioLegend) and analyzed by flow cytometry using AttuneTM NXT.

### In vivo immunogenic cell death evaluations

B16F10 cells were transfected with PCNA-degradation and PCNA-mutant mRNA for 24 h. The treated cells were collected for intratumor injection with the number of 1 ×10^5^ cells per mouse on B16F10 mouse model. Tumor growth was monitored by measuring the length (L) and width (W) of tumors every other two days. The volume (V) of the tumor was analyzed by the formula V = (L x W^2^)/2

### Transcriptome sequencing

A549 cells transfected with control cmRNA or cmRNA encoding PCNA degrader were collected and quickly frozen in liquid nitrogen. Total RNA was extracted and mRNA was purified from totol RNA by poly-T oligo-attached magnetic beads. Then mRNA was fragmented and first strand cDNA was synthesized using random hexamer primer as well as M-MuLV reverse transcriptase. After degrading the RNA by RNaseH, second strand cDNA was synthesized subsequently by DNA polymerase I and dNTP. The library was constructed by the adenylation of 3’ ends and ligation with adaptor containing hairpin structure of cDNA fragments. After qualification of the library, sequencing was performed using Illumina NovaSeq 6000. Clean data with high quality was used for subsequent analysis. Firstly, clean reads were aligned to reference genome by Hisat2 (v2.0.5), then the read numbers mapped to each gene were counted by featureCounts (v1.5.0-p3). Gene expression levels were evaluated by FPKM which was calculated based on the length of the gene and reads count mapped to this gene. Differential expression analysis of two groups (three biological replicates per group) was performed by DESeq2 R package (1.20.0). The resulting P-values were adjusted using Benjamini and Hochberg’s approach for controlling the false discovery rate. Padj<=0.05 and |log2(foldchange)|>=1 was set as the threshold for significantly differential expression. Gene ontology (GO) enrichment analysis as well as KEGG pathway enrichment analysis of differentially expressed genes was implemented by clusterProfiler R package (3.8.1).

### Mouse models

6-8 weeks female C57BL/6 J mice were obtained from Charles Rivers (Shanghai, China). Mice were fed in the Genepharma Experimental animal center (Suzhou, China) and maintained under standard laboratory conditions. The animals were placed in a 12/12 h light/dark cycle at 25±2°C and 50% humidity. The mice were fed with a regular diet and housed and monitored in a pathogen-free environment. Mouse care and experimental procedures were performed with the approval of the animal care committee of Animal Ethics. C57BL/6 J mice were inoculated with B16F10 cells by subcutaneous injection of 5 × 10^5^ cells to the right flank. After 7 days’ tumor inoculation, the tumor volumes reached about 40 mm^3^, and then mice were divided into five groups (n = 8 per group) for further experiments. Invivojet-PEI, PCNA degrader-invivojet-PEI mixture and PCNA mutant-invivojet-PEI mixture were administered intratumorally, either 8, 11, and 14 days following B16F10 inoculation. 50 μg cmRNA is used for one dose, and100 μg PD-L1 were administered intraperitoneally either 7, 10, and 13 days following B16F10 inoculation. Tumor growth was monitored by measuring the length (L) and width (W) of tumors every other two days. The volume (V) of the tumor was analyzed by the formula V = (L x W^2^)/2.

### TUNEL assay

Tumor tissues were fixed in 4% paraformaldehyde for 48 h, then were equilibrated in 20% sucrose solution overnight, and switched to 30% sucrose solution and incubated overnight for cryoprotection. After then tissues were embedded in OCT media and sectioned using a Cryostat, setting for 10 μm per section. Mounted sections were afterwards processed for TUNEL assay. TUNEL assay was performed using TUNEL system kit (KeyGEN BioTECH) according to manufacturer’s recommended protocol.

### GFP degradation in vitro

293T cells were seeded in a 24 well plate at a density of 1.2 ×10^5^ cells per well overnight. Cells were transfected with 400ng GFP-H2B cmRNA per well on the second day. After 6 hours, transfected with 500ng GFP-DASP cmRNA per well for 24 h to 48 h, meanwhile treated with 10 μM or 20 μM MG132 (Sigma). Fluorescent images were taken with Olympus microscope. The average fluorescent intensity was analyzed by flow cytometry using AttuneTM NXT.

### Statistical analysis

GraphPad Prism V.6.01 software was used for statistical analysis. In vivo data were expressed as mean ±standard errors of means (SEMs). In vitro data are presented as the mean ± SEMs from three independent experiments performed at least in duplicate. An independent t-test was used for comparisons of two treatment conditions. In the comparisons of three or more treatment conditions, an ANOVA with Tukey’s post-hoc test was used. Statistical significance was accepted for p values < 0.05.

## References

1. K. M. Sakamoto, et al., Protacs: Chimeric molecules that target proteins to the Skp1-Cullin-F box complex for ubiquitination and degradation. Proc Natl Acad Sci U S A 98, 8554–8559 (2001).

2. M. Bekes, D. R. Langley, C. M. Crews, PROTAC targeted protein degraders: The past is prologue. Nat. Rev. Drug Discov. 21, 181–200 (2022).

3. M. Pettersson, C. M. Crews, PROteolysis TArgeting Chimeras (PROTACs) - Past, present and future. Drug Discov Today Technol 31, 15–27 (2019).

4. J. Liu, et al., TF-PROTACs enable targeted degradation of transcription factors. J. Am. Chem. Soc. 143, 8902–8910 (2021).

5. S. An, L. Fu, Small-molecule PROTACs: An emerging and promising approach for the development of targeted therapy drugs. EBioMedicine 36, 553–562 (2018).

6. G. M. Burslem, C. M. Crews, Proteolysis-Targeting chimeras as therapeutics and tools for biological discovery. Cell 181, 102–114 (2020).

7. S. Lim, et al., BioPROTACs as versatile modulators of intracellular therapeutic targets including proliferating cell nuclear antigen (PCNA). Proc Natl Acad Sci U S A 117, 5791–5800 (2020).

8. B. Baptista, R. Carapito, N. Laroui, C. Pichon, F. Sousa, MRNA, a revolution in biomedicine. Pharmaceutics 13 (2021).

9. J. Yang, et al., Intratumoral delivered novel circular mRNA encoding cytokines for immune modulation and cancer therapy (2021).

10. G. Kroemer, C. Galassi, L. Zitvogel, L. Galluzzi, Immunogenic cell stress and death. Nat. Immunol. 23, 487–500 (2022).

11. A. Ahmed, S. Tait, Targeting immunogenic cell death in cancer. Mol Oncol 14, 2994–3006 (2020).

12. T. Ye, et al., Chemokine CCL17 affects local immune infiltration characteristics and early prognosis value of lung adenocarcinoma. Front Cell Dev Biol 10, 816927 (2022).

13. S. Stutte, et al., Requirement of CCL17 for CCR7-and CXCR4-dependent migration of cutaneous dendritic cells. Proc Natl Acad Sci U S A 107, 8736–8741 (2010).

14. E. Martinenaite, et al., CCL22-specific T Cells: Modulating the immunosuppressive tumor microenvironment. Oncoimmunology 5, e1238541 (2016).

15. M. Rapp, et al., CCL22 controls immunity by promoting regulatory T cell communication with dendritic cells in lymph nodes. J. Exp. Med. 216, 1170–1181 (2019).

16. M. Moossavi, N. Parsamanesh, A. Bahrami, S. L. Atkin, A. Sahebkar, Role of the NLRP3 inflammasome in cancer. Mol. Cancer 17, 158 (2018).

17. L. Xu, W. Yu, H. Xiao, K. Lin, BIRC5 is a prognostic biomarker associated with tumor immune cell infiltration. Sci Rep 11, 390 (2021).

18. A. Wadhwa, A. Aljabbari, A. Lokras, C. Foged, A. Thakur, Opportunities and challenges in the delivery of mRNA-based vaccines. Pharmaceutics 12 (2020).

19. M. Cardano, C. Tribioli, E. Prosperi, Targeting proliferating cell nuclear antigen (PCNA) as an effective strategy to inhibit tumor cell proliferation. Curr Cancer Drug Targets 20, 240–252 (2020).

20. S. M. Banik, et al., Lysosome-targeting chimaeras for degradation of extracellular proteins. Nature 584, 291–297 (2020).

21. Z. Li, et al., Allele-selective lowering of mutant HTT protein by HTT-LC3 linker compounds. Nature 575, 203–209 (2019).

22. D. Takahashi, et al., AUTACs: Cargo-Specific degraders using selective autophagy. Mol. Cell 76, 797–810 (2019).

23. S. Lim, et al., BioPROTACs as versatile modulators of intracellular therapeutic targets including proliferating cell nuclear antigen (PCNA). Proceedings of the National Academy of Sciences 117, 5791–5800 (2020).

24. M. F. Chretien, A. Chassevent, K. Malkani, A. Rebel, Flow cytometric DNA analysis in the diagnosis of lung tumors. A comparison with conventional methods. Anal Quant Cytol Histol 10, 251–255 (1988).

25. M. Mathew, T. Enzler, C. A. Shu, N. A. Rizvi, Combining chemotherapy with PD-1 blockade in NSCLC. Pharmacol Ther 186, 130–137 (2018).

26. J. Humeau, S. Levesque, G. Kroemer, J. G. Pol, Gold standard assessment of immunogenic cell death in oncological mouse models. Methods Mol Biol 1884, 297–315 (2019).

27. A. Viola, A. Sarukhan, V. Bronte, B. Molon, The pros and cons of chemokines in tumor immunology. Trends Immunol. 33, 496–504 (2012).

28. K. V. Swanson, M. Deng, J. P. Ting, The NLRP3 inflammasome: Molecular activation and regulation to therapeutics. Nat. Rev. Immunol. 19, 477–489 (2019).

29. Y. Huang, W. Xu, R. Zhou, NLRP3 inflammasome activation and cell death. Cell. Mol. Immunol. 18, 2114–2127 (2021).

30. Y. N. Lamb, BNT162b2 mRNA COVID-19 vaccine: First approval. Drugs 81, 495–501 (2021).

31. W. Xie, B. Chen, J. Wong, Publisher Correction: Evolution of the market for mRNA technology. Nat. Rev. Drug Discov. 20, 880 (2021).

32. J. D. Gillmore, et al., CRISPR-Cas9 in vivo gene editing for transthyretin amyloidosis. N Engl J Med 385, 493–502 (2021).

33. Q. Cheng, et al., Selective organ targeting (SORT) nanoparticles for tissue-specific mRNA delivery and CRISPR-Cas gene editing. Nat. Nanotechnol. 15, 313–320 (2020).

34. U. Sahin, et al., An RNA vaccine drives immunity in checkpoint-inhibitor-treated melanoma. Nature 585, 107–112 (2020).

